# Safety processing shifts from hippocampal to network engagement across adolescence

**DOI:** 10.1101/2025.06.23.661189

**Authors:** Yubing Zhang, Madeline Coates, Marta I. Garrido, Sarah M. Tashjian

## Abstract

Adolescence is a critical period that requires balancing exploration of uncertain and novel environments while maintaining safety. This task requires sophisticated neural integration of threat and safety cues to guide behavior. Yet little work has been conducted on threat and safety processing outside of conditioning paradigms, which, while valuable, lack the complexity to identify how the adolescent brain supports distinguishing threat from safety when both are present and as task contingencies change. In the current study, we take an approach that expands on elements of differential conditioning as well as conditioned inhibition. We recorded brain responses to external threat and self-oriented protection cues to examine how the adolescent brain supports threat-safety discrimination using 7-Tesla functional magnetic resonance imaging (fMRI). Our findings reveal an adolescent transition in the neural mechanisms supporting accurate threat-safety discrimination, with younger adolescents (12-14 years) relying predominantly on the hippocampus and older adolescents (15-17 years) utilizing a more integrated circuit involving the hippocampus and anterior ventromedial prefrontal cortex (vmPFC) connectivity. Our results clarify how competition between threat and safety cues is resolved within the visual cortex, demonstrating enhanced perceptual sensitivity to protection that is independent of threat. By examining the dynamic encoding of safety to different stimuli, the current study advances our understanding of adolescent neurodevelopment and provides valuable insights into threat-safety discrimination beyond conventional conditioning models.

**Highlights:**

- Protection is more strongly weighted than threat in adolescent safety estimation.
- Hippocampus aids accurate safety detection in younger adolescents.
- Hippocampal-vmPFC connectivity aids accurate safety detection in older adolescents.
- Protection enhances visual processing, reflecting perceptual prioritization.

## 1. Introduction

Identifying threat is critical for survival, yet overestimations of threat can prevent individuals from accomplishing other goals, such as exploring and learning. A delicate balance between threat and safety can be difficult to achieve, particularly during adolescence (Bach & Dayan, 2017; LeDoux & Pine, 2016). This crucial developmental period demands exploring uncertain environments to gain independence and experience (Crone & Dahl, 2012; Gopnik et al., 2017; Murty et al., 2016; Somerville et al., 2017; Spear, 2000) while navigating neurobiological changes that can impede threat-safety discrimination (Larsen & Luna, 2018; Hartley & Somerville, 2015; Towner et al., 2023). Inability to accurately identify threat and safety can contribute to harmful risk-taking behaviors during adolescence, like reckless driving and substance abuse (Hartley & Somerville, 2015; Galván & Rahdar, 2013). Also relevant is the onset of mental health difficulties in adolescence, such as anxiety, characterized by aberrant threat and safety processing (Casey et al., 2008; Ciranka & van den Bos, 2021; Lee et al., 2014). However, little work has been conducted on threat-safety discrimination during development outside of conditioning paradigms (Fullana et al., 2020; Odriozola & Gee, 2021). In the current study, we adopt Tashjian and colleagues’ (2025) multi-component definition of safety, which frames safety estimation as emerging from the dynamic integration of external threat assessment and self-oriented protective evaluation. This process involves coordinated processing between memory systems that encode threat-relevant experiences (hippocampus) and executive systems that evaluate protective capabilities and regulate threat responses (vmPFC). We also employ 7-Tesla functional magnetic resonance imaging (7T fMRI) to achieve unprecedented resolution for identifying related neural systems. This approach uniquely captures how the maturing adolescent brain balances sensory signals with emerging metacognitive abilities (Weil et al., 2013), revealing developmental mechanisms for safety estimation beyond conventional conditioning models.

Safety has not been consistently and formally defined in the existing literature, and it remains poorly understood compared to threat (Laing & Harrison, 2021). The most frequently used experimental paradigms for investigating safety build on fear conditioning models and adopt different operationalizations of safety (Grasser & Jovanovic, 2021; Sangha et al., 2020). In differential conditioning and extinction learning, safety is conceptualized as an ambiguous state characterized by the absence of threat, whereas in conditioned inhibition and conditioned discrimination, safety involves the learned suppression of responses to threat cues (Laing & Harrison, 2021; Laing et al., 2022). One limitation of these approaches is that they focus on external threat signals without a clear demonstration of how safety fluctuates independent of threat. In real-world settings, safety estimation often requires a distinct process of recognizing certain cues or contexts that signal safety even in the presence of threats. Safety, conceptualized as a dynamic integration of threat and protection in the current study, is most closely related to conditioned inhibition. However, while conditioned inhibition entails learning inhibitory cues signaling no aversive outcome, our study requires individuals to understand that self-oriented protection actively mitigates the existing threat to different degrees and therefore creates a “meta-representation” of safety (Tashjian et al., 2025). This evaluation process may involve integrating multiple functions, including threat-processing, self-representation, and metacognitive assessment (Kveraga et al., 2007; Tashjian et al., 2021). Importantly, understanding the resources available to support self-related processing is still developing during adolescence, potentially contributing to difficulties in threat-safety discrimination that have been overlooked in conditioning paradigms. To our knowledge, the current study is the first to examine self-oriented protective components of safety processing in human adolescents.

A range of animal studies and human fMRI studies support the notion that threat and protection evaluations are computed in distinct neural networks (Kong et al., 2014; Tashjian et al., 2025; Wen et al., 2024). Specifically, the posterior ventromedial prefrontal cortex (vmPFC), along with the defensive circuit (including the periaqueductal gray, amygdala, insula, and hypothalamus), is involved in detecting and responding to potential threats (Greco & Liberzon, 2016; Mobbs et al., 2020; Tovote et al., 2016). In terms of safety, the anterior vmPFC has been identified as a critical region for the retention of extinction, identification of self-oriented safety cues, and inhibition of stress-induced behavioral and psychological responses (Battaglia et al., 2022; Harrison et al., 2017; Maier et al., 2006; Phelps et al., 2004; Schiller et al., 2008; Tashjian et al., 2021; Tashjian et al., 2025). The hippocampus facilitates safety estimation by encoding memories of successfully confronted threatening situations and supporting strategic decisions in safe states (Micale et al., 2017; Qi et al., 2018). Additionally, the anterior cingulate cortex (ACC) monitors conflict between original threat and new safety information, mediating behavioral flexibility (Jovanovic et al., 2013; Tanida et al., 2018; Tashjian et al., 2025; Wu et al., 2023). While the distributed neural architecture for threat-safety processing is well-documented in adults, research indicates that these regions follow different developmental trajectories during adolescence. In particular, the hippocampus and ACC mature earlier than the vmPFC (Casey et al., 2016; Fuster, 2002; Lau et al., 2011; Mills et al., 2014), creating unique imbalances that may influence adolescent safety evaluation. Given these neurodevelopmental patterns, examining safety processing through an integrated threat and self-oriented protection framework is essential to understand how developing neural circuits differentially support these complementary dimensions of safety evaluation.

Evidence is mixed regarding how safety estimation develops during adolescence. Some studies reveal a progressive development in extinction and differential conditioning during childhood, which reaches maturity by adolescence (Abend et al., 2020; Britton et al., 2013; Waters et al., 2017; Widegren et al., 2025). These findings conflict with others that report an adolescent-specific reduction in safety detection (Ganella et al., 2018; Lau et al., 2011; Pattwell et al., 2012). Discrepancies and ambiguities in the literature are attributable, in part, to methodological and measurement differences. For instance, current standard 3 Tesla fMRI encounters drop-out and low signal-to-noise resolution in subcortical and midline prefrontal regions (Morris et al., 2019). Many studies index threat-safety discrimination through changes in skin conductance response (SCR), but SCR is poorly suited to fMRI as supine posture and the cold MRI environment attenuate SCR responses and signal quality (Morriss et al., 2019). Additionally, examining adolescents as a group rather than testing age-related changes during adolescence obscures non-linear developmental shifts. These factors widen the existing gap in knowledge regarding developmental changes in higher-order threat-safety discrimination. The current study aims to expand and clarify the understanding by incorporating external threat and self-oriented protection cues that vary in intensity independently and in their dynamic interaction. We hypothesized that: (a) Self-oriented protection is critical to safety evaluation during adolescence and recruits distinct neural systems from those engaged by external threat; (b) Adolescent safety evaluations vary with the intensity and order of threat and safety cues; (c) Because our task requires higher-order cognitive processes such as information integration, we expected age-related differences with younger adolescents demonstrating less accurate evaluations; (d) Although the vmPFC has been identified as a central hub for safety evaluation in adults, it continues to mature throughout adolescence. Therefore, adolescents were expected to rely more on subcortical systems, particularly at younger ages.

## 2. Methods

### 2.1. Participants and Ethics

Thirty-five participants aged 12 to 17 years old were recruited through flyers and online postings. We designed the study to interrogate changes during adolescence, given robust neural and behavioral changes among individuals in this age range and the fact that studies that examine age differences among adolescents are more likely to successfully characterize non-linear developmental processes than those that consider adolescents as a single group compared to adults. Our inclusion criteria were as follows: (1) not claustrophobic; (2) no metal contraindications; (3) not pregnant; (4) no hearing or sight difficulties; (5) no medicated neurological or psychiatric conditions; and (6) fluent in verbal and written English. Two participants were excluded from all analyses: One due to an incomplete MRI session and another due to a potential neurological disorder identified during scanning. This resulted in a final sample of 33 participants (*M_Age_* = 14.88, *SD_Age_* = 1.47, 19 females 58%). All methodology was approved by the University of Melbourne Human Research Ethics Committee (27613). Participation was voluntary, and participants were compensated for their time. Informed assent was obtained from the adolescent, and informed consent from their parent/guardian.

### 2.2. Procedure

Prior to scanning, participants completed an online demographic questionnaire and a series of questionnaires. The MRI session consisted of a structural scan and three fMRI tasks, one of which was the Safety Estimation Task described here. The other two tasks are out of scope and will be described elsewhere. For the Safety Estimation Task, participants received task instructions and completed five practice trials during the structural scan. Participants completed four functional runs of the task (approximately 6 minutes each).

### 2.3. Task Design

The Safety Estimation Task was adapted from prior work examining safety processing in adults (see Tashjian et al., 2025 for task development). Participants were instructed to imagine fictitious battles fighting against threatening animals (cat, goose, lion, or grizzly) using powerful weapons (fist, stick, gun, or grenade). Each cue (animal or weapon) was presented in isolation, and their presentation was counterbalanced. Thus, the first cue was presented without knowledge of the second cue. The second cue completed the information for the given trial. After both cues were presented, participants saw the win/lose outcome for 2 seconds. Outcomes were paired with either a loud, unpleasant white noise (loss) or no noise (win). Outcomes were unrelated to participant choices and instead were predetermined based on experimentally established probabilities for the weapon-animal pair presented. As shown in the safety continuum (Fig. 1b), the safety probabilities were designed to be identically balanced, such that participants had the same likelihood of winning (50% chance) whether equipped with the most powerful weapon while facing the most threatening animal or equipped with the least powerful weapon while facing the least threatening animal. Participants were asked to make binary judgments about whether they thought they would win or lose the battle against each animal using the weapon provided. Judgments were made twice on each trial, in response to the first and second cue presentations (Fig. 1a). Each weapon/animal cue was shown for 6 seconds maximum (offset to participant button press), followed by a jittered .5 to 2-second interstimulus interval (ISI). Trials were separated by an inter-trial interval (ITI), presented for .5 to 2 seconds with duration jittered. All participants completed 160 trials, with each animal-weapon pair presented 10 times. The task was programmed using PsychoPy v2023.1.2.

**Fig. 1.**
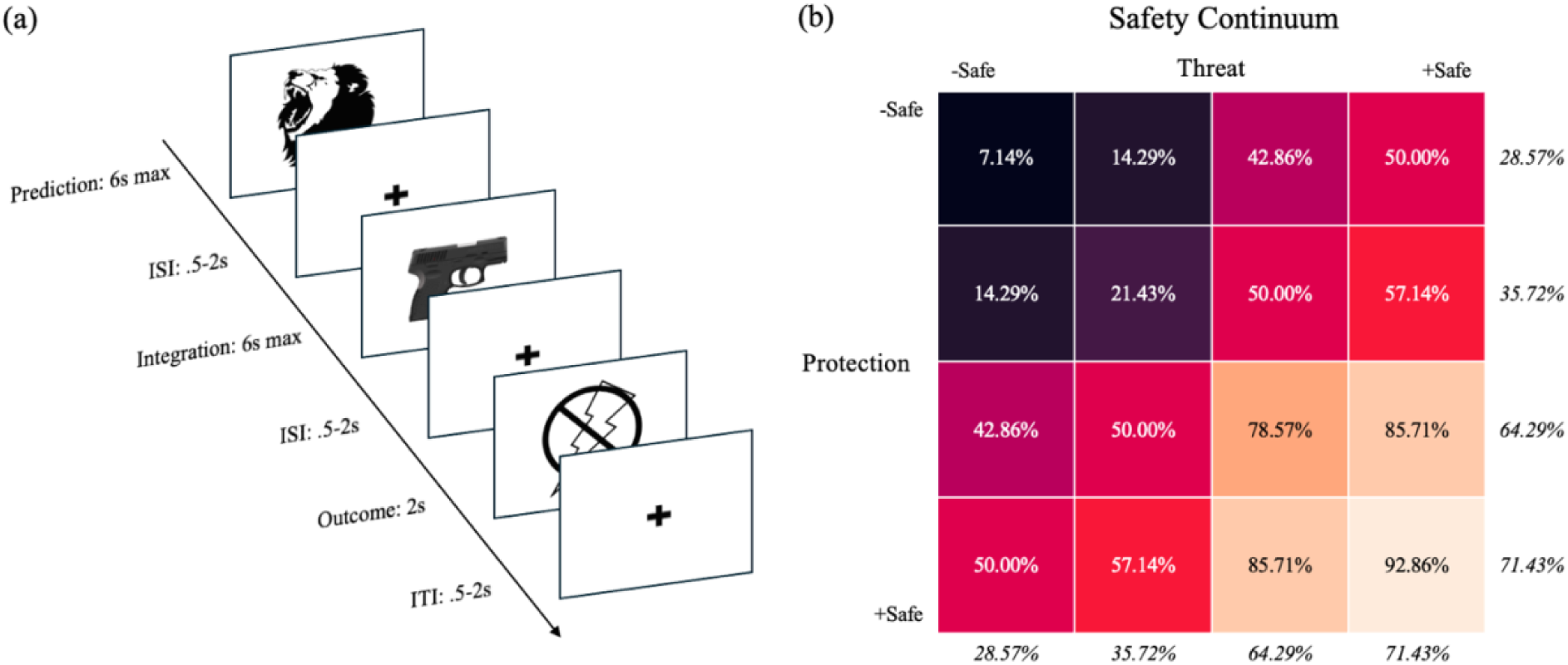
(a) An example trial of the Safety Estimation Task. Participants saw an animal-weapon pair with each cue presented in isolation and in counterbalanced order. Participants were instructed to make binary choices about whether they thought they would win or lose the battle in response to the first and second cue presentations. Participants saw the outcome of the battle for 2 seconds, which was paired with either a loud, unpleasant white noise (loss) or no noise (win). (b) The predetermined experimental safety continuum showing the probability of winning for each condition (protection continuum order: fist, stick, gun, grenade; threat continuum order: cat, goose, lion, grizzly). Values inside each grid cell indicate the probability taking into account the animal-weapon pairing, while values in italics indicate the average winning probability for each cue across all pairings. The safety continuum was designed to be identically balanced. For example, participants had a 50% chance of winning with a combination of the most powerful weapon and the most threatening animal, which was identical to the probability with a combination of the least powerful weapon and the least threatening animal. Similarly, the least powerful weapon (fist, 28.57%) had an identical winning probability as the most threatening animal (grizzly, 28.57%), and the same for the most powerful weapon (grenade, 71.43%) compared to the least threatening animal (cat, 71.43%).

### 2.4. MRI Data Acquisition

Structural and functional MRI data were acquired at the Melbourne Brain Centre Imaging Unit (MBCIU) using a Siemens 7T Plus scanner equipped with a 32-channel head coil. Structural (T1-weighted) images were obtained using a magnetization-prepared 2 rapid acquisition gradient echo sequence (MP2RAGE; Marques et al., 2010) (TR = 5000 ms, TE = 2.04 ms, flip angle 1 = 4 deg, flip angle 2 = 5 deg, field of view = 240 mm, slice thickness = 0.75 mm). Functional (T2-weighted) images were collected using a gradient echo-planar imaging sequence (EPI; Moeller et al., 2010): TR = 800 ms, TE = 22.20 ms, flip angle = 45 deg, field of view = 208 mm, slice thickness = 1.60 mm. A total of 84 interleaved slices were acquired parallel to the anterior-posterior commissure line with a multiband acceleration factor of 6.

### 2.5. Data Analysis

We separately examined participants’ judgments in response to the first and second cues. The first judgment, safety prediction, reflects how participants discriminated between threat and safety based on the type of cue (animal versus weapon). The second judgment, safety integration, reflects how participants integrated information from both cues (animal and weapon combined). All cues were presented an equal number of times in both first and second positions. We calculated participants’ task accuracy for each condition as the difference between the proportion of trials they predicted as wins and the experimentally established probability of winning.

MRI data were pre-processed using fMRIPrep 23.2.1 (Esteban et al., 2019) and analyzed using FSL (Jenkinson et al., 2012). Structural scans were corrected for intensity non-uniformity, skull stripped, and normalized to the ICBM 152 Nonlinear Asymmetrical template (2009c) through nonlinear registration. Functional scans underwent motion correction (24 parameters including 6 standard and 18 extended derivatives), susceptibility distortion correction, and co-registration to the T1-weighted reference image using boundary-based registration. A first-level general linear model (GLM) was defined for each run of the Safety Estimation Task with 11 regressors modeling different trial components. Regressors 1-7 modeled cue presentations: (1) Threat First (animal presented as the first image), (2) Protection First (weapon presented as the first image), (3) Threat Second, (4) Protection Second, (5) Win Outcome, (6) Lose Outcome, (7) Fixation (ITI + ISI). Four additional regressors were included for cue presentations (regressors 1-4), consisting of pre-determined win/lose probability as parametric modulators (Fig. 1b). Both standard and parametric regressors were modeled with a canonical double-gamma hemodynamic response function (HRF) for a duration from image onset to offset. Parametric modulation regressors were orthogonalized with respect to the lower-order regressors (Mumford et al., 2015). Temporal derivatives were included for all regressors to reduce slice-timing differences and variability in the HRF delays across regions, thereby enhancing model fit. The four runs of the Safety Estimation Task were combined for each participant using a fixed-effect voxel-wise second-level model in FEAT. Group-level analyses were performed using the FMRIB Local Analysis of Mixed Effects (FLAME1) module in FSL (Beckmann et al., 2003). Z-statistic images were thresholded using a cluster-forming threshold of *Z* > 3.1 and a familywise error-corrected cluster significance threshold of *p* < .05 based on the Theory of Gaussian Random Fields (Poline et al., 1997; Worsley, 2001). Statistical maps of all analyses were projected onto a standard MNI brain. Group activation maps were visualized using MRIcroGL 14.3.1 (https://www.nitrc.org/projects/mricrogl/). We further assessed the interaction between condition (threat or safety) and presentation order (first or second) using a separate GLM with four regressors: (1) Condition, (2) Order, (3) Interaction, and (4) Intercept, while keeping all other modeling steps identical as above.

Based on the GLM results indicating increased left hippocampal activation for protection relative to threat as the first cue (Fig. 3b), we conducted ROI analyses for further exploration. We used a ventral hippocampus mask defined from a probabilistic atlas of the medial temporal lobe, independent from the current fMRI data (Fig. 3a; Hindy & Turk-Browne, 2016; Meyer et al., 2019). For each participant, mean activation within the left hippocampus for the Protection First > Threat First contrast was extracted from the filtered 4D data at the group level using FSLMEANTS and was analyzed in R. A generalized psychophysiological interaction analysis (gPPI; Friston et al., 1997) was also performed to investigate task-related modulation of functional connectivity patterns between the hippocampus and other brain regions identified in prior work as relevant for safety processing, specifically the anterior vmPFC and ACC (Meyer et al., 2019; Tashjian et al., 2025). The same ventral hippocampus ROI from the activation analysis was used. The standard-space mask was transformed into subject space using FLIRT, and the average time series of all voxels within the mask was extracted using FSLMEANTS. The product between the hippocampus time series (physical regressors) and individual regressors for each condition (psychological regressors) was included in each participant’s first-level GLM design matrix. The psychological regressors were zero-centered, and the physical regressors were demeaned. Functional connectivity estimates for the anterior vmPFC (“frontal medial cortex”) and the caudodorsal subregion of the ACC were then extracted from the Protection First condition and further analyzed in R. These regions were selected based on prior findings of safety circuitry in adults (Meyer et al., 2019; Tashjian et al., 2025). Anterior vmPFC and ACC masks were defined according to the Harvard-Oxford cortical and subcortical structural atlases, independent from the current fMRI data (Fig. 3a).

## 3. Results

### 3.1. Behavioral Results

Participants estimated a greater likelihood of winning when exposed to cues with higher experimentally established safety probabilities (Fig. 2a, 2c), indicating they were able to track safety probabilities successfully. For safety prediction, the source of information affected initial evaluations of the first cue, such that participants estimated a higher probability of winning in response to protection cues compared to threat cues (Fig. 2b). Participants also integrated information from both cues differently based on the order of presentation and the changes in safety probability (Fig. 2d). Specifically, they were more likely to update their safety judgments when threat cues were presented first and followed by protection cues, and more likely to stick to the initial decisions in opposite conditions (*β* = .36, *SE* = .06, *z* = 6.02, *p* = 1.72 x 10^-09^). For example, if participants first saw a lion, they were more likely to revise their safety judgment when receiving a grenade as their weapon, and more likely to keep their safety judgment the same if they first received the grenade and subsequently saw they were battling a lion. Participants were also more likely to update their judgments when safety probabilities were increased from the first cue to the second cue, compared to when safety probabilities were decreased (*β* = .96, *SE* = .17, *z* = 5.64, *p* = 1.52 x 10^-08^). Despite equivalent experimentally established win/lose probabilities, participants judged their winning likelihood as higher when they had powerful weapons and were battling threatening animals, compared to when they had weaker weapons and were battling less threatening animals (Fig. 2e-f). This asymmetric weighting is evidenced by participants’ tendency to judge scenarios as safer when protection was strong (powerful weapons against threatening animals) compared to when threat was weak (weak weapons against non-threatening animals), despite these conditions having identical win probabilities. Together, these results indicate that, compared to threat cues, protection cues were more strongly weighted in both initial safety predictions and subsequent safety integrations.

**Fig. 2.**
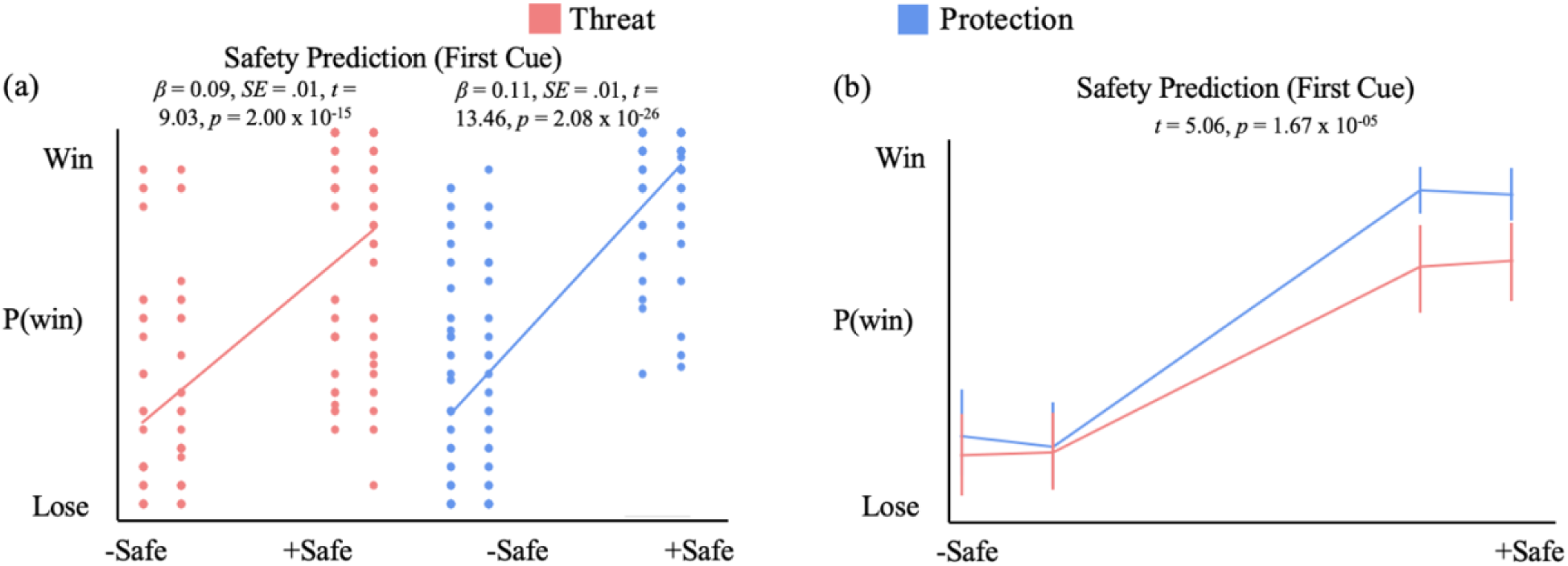
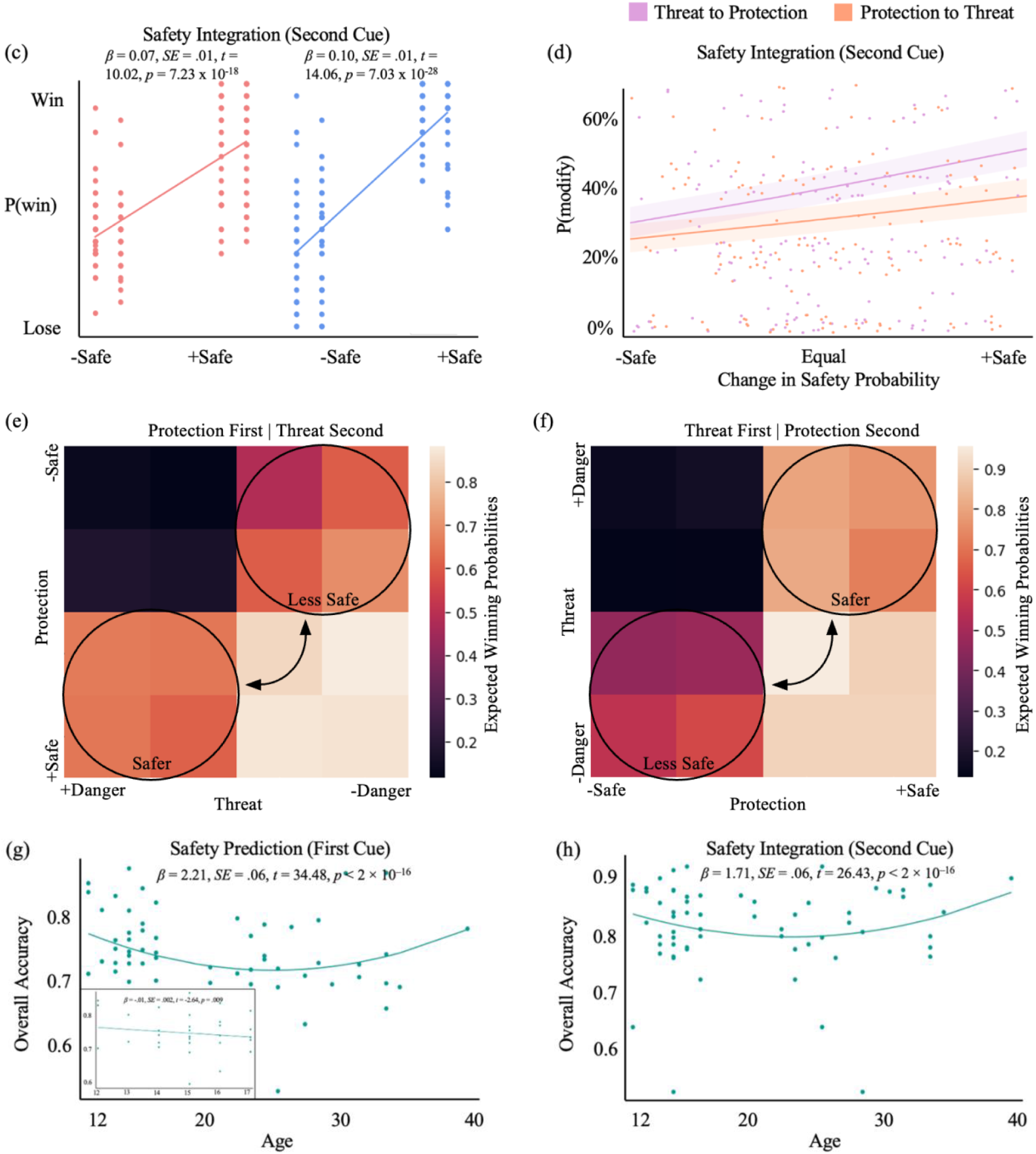
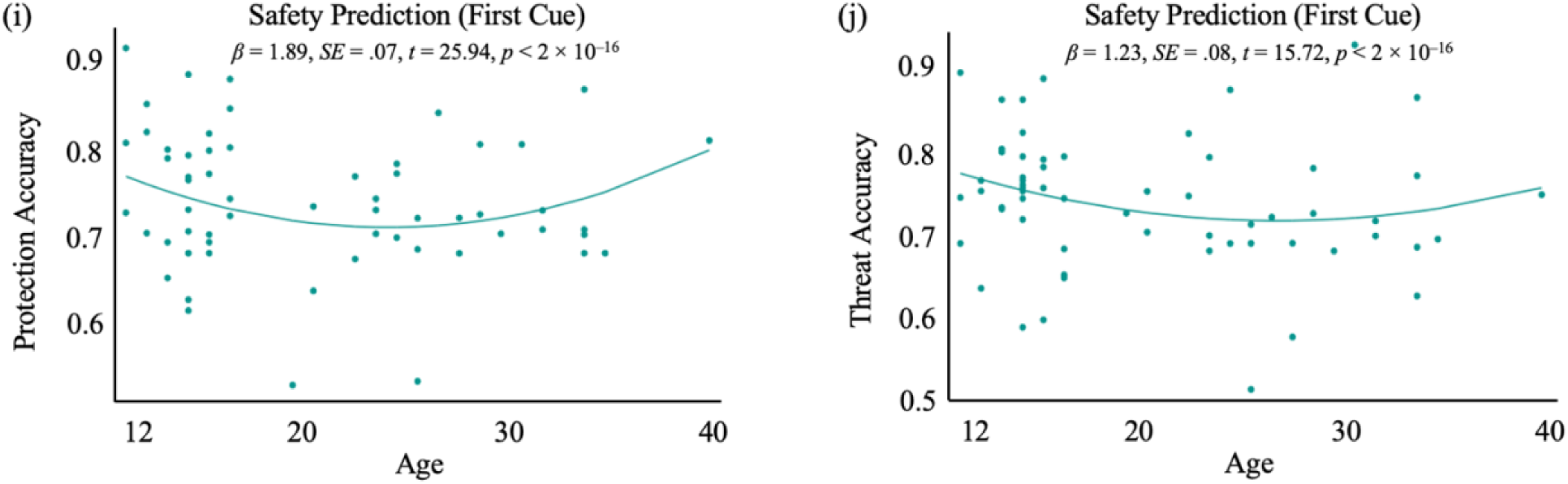
Mixed effects logistic regression showing the relationship between experimentally established safety probabilities and participants’ safety judgments. Results suggested that participants were able to track the task safety contingencies successfully for the first (a) and second (c) cues. (b) For safety prediction, participants estimated a higher probability of winning when protection was presented as the first cue. (d) For safety integration, participants were more likely to update their safety evaluations when threat cues were followed by protection cues and when safety probabilities were increased. (e-f) Expected winning probabilities. Participants expected higher probabilities of winning when they had powerful weapons and were faced with threatening animals, compared to when they had less powerful weapons and were faced with weak animals, regardless of the order of presentation. Quadratic associations were observed between age and the overall accuracy for safety prediction (g) and safety integration (h) in the full sample, with the accuracy decreasing during adolescence and increasing from the mid-twenties. The inset plot in (g) further shows that age was negatively associated with participants’ overall accuracy for safety prediction when considering the adolescent sample only. For safety prediction, quadratic associations were also observed between age and the accuracy of both protection (i) and threat (j), with the accuracy decreasing during adolescence and increasing from the mid-twenties. All results remain the same after removing outliers at +2SD.

In our adolescent fMRI sample, age was negatively associated with overall accuracy during safety prediction (Fig. 2g). When extending our inquiry to include a separate adult sample (sourced from Tashjian et al., 2025 in which the same Safety Estimation Task was used; adults: *N* = 30, *M*_age_ = 27.83, *SD*_age_ = 4.86, range 20-40 years, 15 females 50%; full sample: *N* = 63, *M*_age_ = 24.18, *SD*_age_ = 7.23, range 12-40 years, 34 females 54%), a quadratic association was observed between age and overall accuracy (Fig. 2g). This clarified the developmental pattern, revealing that both younger adolescents and adults tended to make more accurate safety judgments than older adolescents. When separately assessed, protection and threat displayed similar quadratic trends during safety prediction in the full sample (Fig. 2i-j). In terms of the integration of threat and protection cues, age was not significantly associated with accuracy in the adolescent sample (*β* = -.001, *SE* = .003, *z* = -.47, *p* = .640), but again, a quadratic association was observed in the full sample (Fig. 2h).

### 3.2. MRI Results

In the current study we used 7T fMRI, which differed in ability to detect subcortical activation compared with Tashjian et al. (2025), which used 3T fMRI. Taking into account potential comparability issues, all MRI results reported below include only the adolescent sample aged 12-17 years. When compared with threat, protection evoked greater activation in bilateral visual processing regions (the temporal fusiform cortex, occipital fusiform gyrus, lateral occipital cortex, and occipital pole), and left motor sensory cortices (precentral and postcentral gyri; Fig. 3b) when considering the first cue. Interestingly, protection also evoked activation in the left hippocampus, which has been previously linked to safety estimation (Meyer et al., 2019). Greater activation was elicited in the right lateral occipital cortex for threat compared to protection as the first cue (Fig. 3c). For the second cue, protection compared to threat elicited greater activation in visual processing regions (the left temporal fusiform cortex, left occipital fusiform gyrus, left lateral occipital cortex, and bilateral occipital pole) and the left postcentral gyrus (Fig. 3d). No voxels survived cluster correction at *Z* > 3.1, *p* < .05 for threat compared to protection as the second cue. No significant clusters survived multiple comparison correction at the whole brain level for the interaction between cue type and presentation order.

**Fig. 3.**
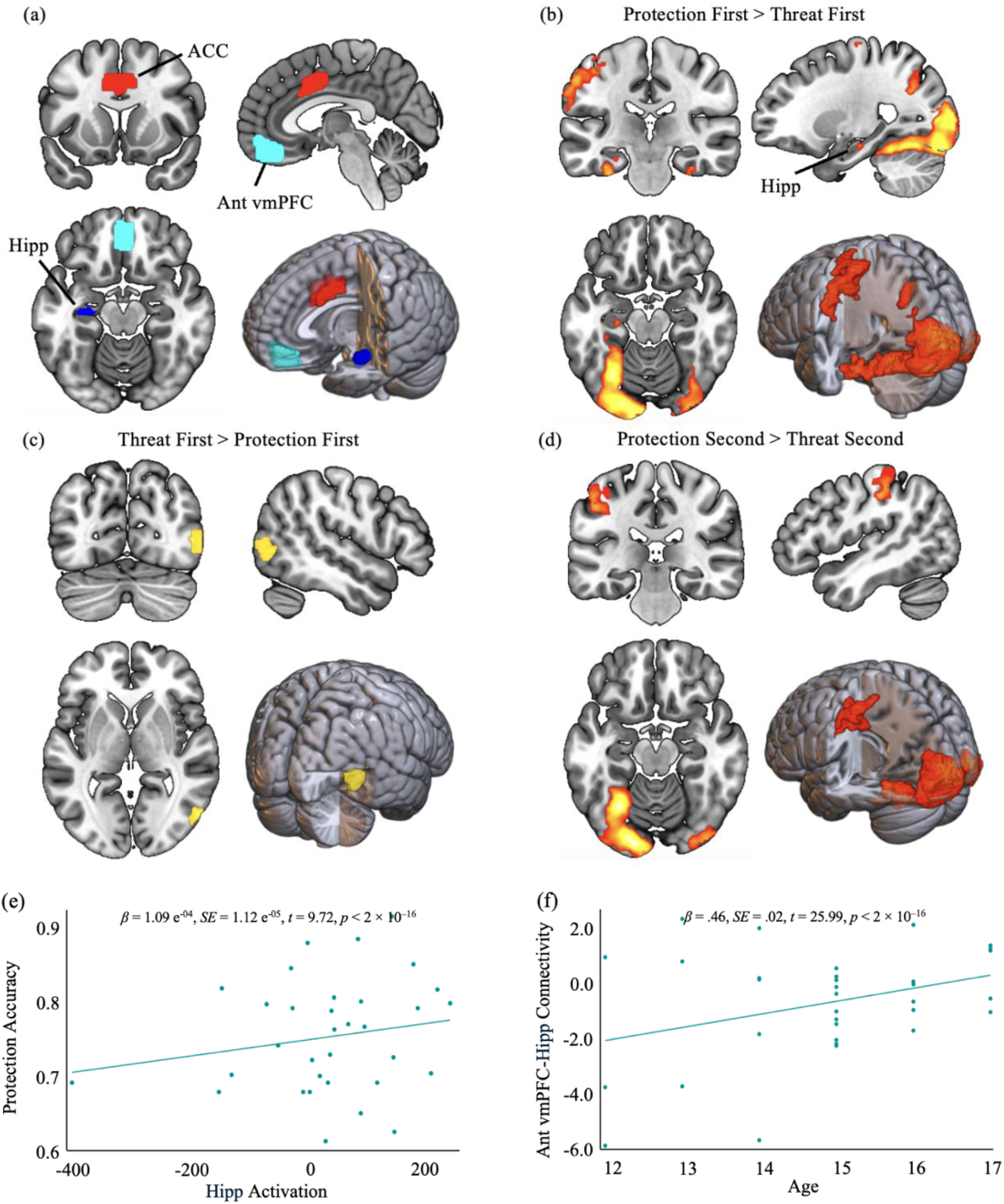
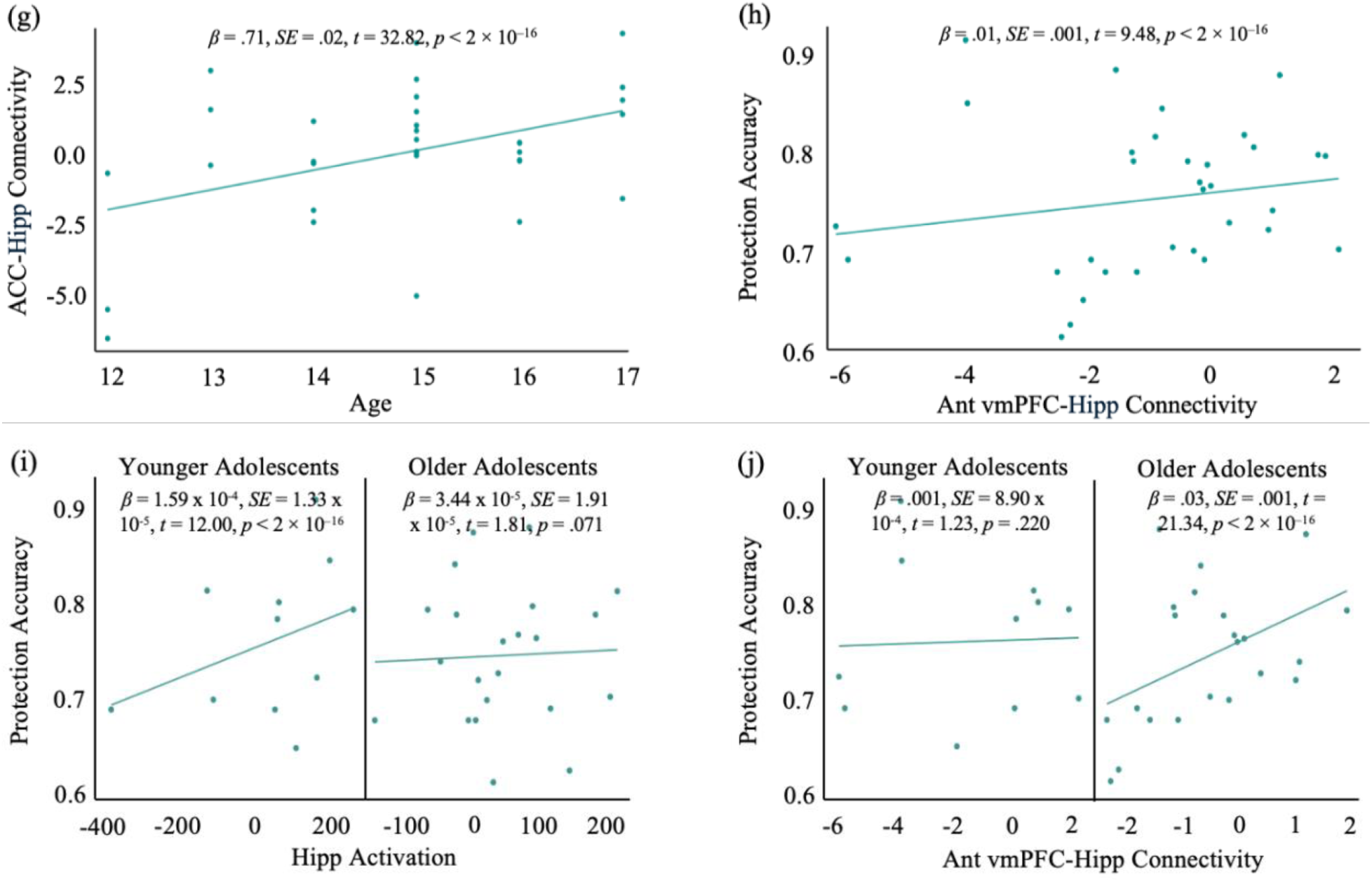
(a) ROIs used in activation and generalized psychophysiological interaction (gPPI) analyses. A ventral hippocampus mask was defined from a probabilistic atlas of the medial temporal lobe; anterior vmPFC and ACC masks were defined according to the Harvard-Oxford cortical and subcortical structural atlases. All masks were independent from the current fMRI data. (b-d) Visualization of significant activation for the contrasts of Protection First > Threat First, Threat First > Protection First, and Protection Second > Threat Second. FLAME1, Z > 3.1, FWE-corrected p < .05. (e) Accuracy for protection cues during the first cue presentation increased with hippocampal activation. (f-g) Hippocampal-ACC and hippocampal-anterior vmPFC connectivity for the Protection First > Threat First contrast increased with age. (h) Accuracy for protection cues during the first cue presentation increased with hippocampal-anterior vmPFC connectivity. (i-j) After using median split to divide adolescents into younger (12-14 years) and older (15-17 years) subgroups, we found that hippocampal activation was positively associated with participants’ accuracy in detecting protection as the first cue in younger but not older adolescents, and hippocampus-vmPFC connectivity was positively associated with participants’ accuracy in older but not younger adolescents. All results remain the same after removing outliers at +2SD. ACC = anterior cingulate cortex; Ant vmPFC = anterior ventromedial prefrontal cortex; Hipp = ventral hippocampus.

We found the increased activation in the left ventral hippocampus for protection compared to threat during the first cue presentation to be particularly intriguing. To further probe this effect, we conducted exploratory ROI and gPPI analyses. ROI analysis revealed that hippocampal activation was not significantly associated with age (*β* = -3.90, *SE* = 15.67, *t* = -.25, *p* = .805). Hippocampal activation when evaluating protection versus threat in response to the first cue was positively associated with participants’ accuracy in estimating protection cues (Fig. 3e). For gPPI analysis, we examined connectivity between the hippocampus and anterior vmPFC as well as the hippocampus and ACC. Connectivity for both ROI pairs was positively associated with continuous age in the adolescent sample (12-17 years) (Fig. 3f-g). A positive association was observed between hippocampal-anterior vmPFC connectivity and participants’ accuracy in the detection of protection during the first cue presentation (Fig. 3h). No significant association was found between hippocampal-ACC connectivity and protection accuracy (*β* = 1.53 x 10^-4^, *SE* = 5.91 x 10^-4^, *t* = .26, *p* = .795). To further probe age effects in hippocampal activation and hippocampal connectivity, we used a median split to examine adolescent subgroups comprised of younger (12-14 years) and older (15-17 years) adolescents. We found that hippocampal activation was positively associated with participants’ accuracy in detecting protection as the first cue in younger but not older adolescents (Fig. 3i). In contrast, hippocampus-vmPFC connectivity was positively associated with participants’ accuracy in older but not younger adolescents (Fig. 3j).

## 4. Discussion

The current study reveals a developmental trajectory in the neural mechanisms supporting threat-safety discrimination. We identified compensatory mechanisms recruited during adolescence, with younger adolescents (12-14 years) relying predominantly on subcortical systems, namely the hippocampus, to accurately distinguish cues, and engagement of subcortical-cortical circuits with increasing age. For older adolescents (15-17 years), connectivity between the hippocampus and anterior vmPFC aided accurate threat-safety discrimination. These findings provide novel insights into a neurodevelopmental shift in safety processing that may have implications for understanding adolescent risk-taking behavior and vulnerability to anxiety (Casey et al., 2008; Galván & Rahdar, 2013; Zacharek et al., 2021). We also found increased engagement of the visual processing regions for the detection of protection compared to threat, reflecting enhanced perceptual sensitivity to salient protection cues that are independent of threat cues, and regardless of age.

We employed a novel paradigm to investigate safety estimation by combining external threat cues with self-oriented protection cues and examining their dynamic interplay. Our design shares key features with traditional paradigms for evaluating safety from the fear conditioning framework, in particular conditioned inhibition and differential conditioning. When presented in isolation during the first half of each trial, threat and protection cues function analogously to CS+ and CS- in differential conditioning, each predicting a different outcome. In trials where a threat cue is immediately followed by a protection cue, protection serves as a partial conditioned inhibitor, signaling a reduced probability of an aversive outcome. While grounded in the standard conditioning framework, our design extends it by implementing probabilistic cue-outcome contingencies to modulate safety and treating protection signals as independent entities, rather than mere inverses of threat cues. This approach better approximates real-world situations where threat and protection often co-occur, and provides a more nuanced understanding of adolescent safety processing. Our results also suggest that adolescents actively tracked the safety probabilities associated with both cue types. We found evidence that threat and protection cues contributed differently to adolescents’ safety perception, with protection having a larger impact on safety judgments.

We observed age-related differences in safety evaluation accuracy, with linear decreases in our adolescent sample and quadratic effects when combining our adolescent sample with a separate sample of adults. The U-shaped pattern suggests that mid-to-late adolescence is a period of vulnerability in threat-safety discrimination. Although recent evidence suggests a progressive development in fear extinction and differential conditioning (Abend et al., 2020; Widegren et al., 2025), our study demonstrates that when examining more complex aspects of threat-safety discrimination—particularly those involving self-oriented factors—adolescents show distinct behavioral patterns. Our neuroimaging results clarify this finding, showing differential mechanisms aiding safety evaluation accuracy. For younger adolescents, accuracy for detecting protection was positively associated with hippocampal activation but not with connectivity between the hippocampus and ACC or vmPFC. Older adolescents displayed the opposite pattern: those with higher hippocampal-anterior vmPFC connectivity demonstrated higher accuracy for detecting protection cues, while hippocampal activation alone no longer predicted their performance. These results demonstrate a fundamental developmental shift in neural mechanisms supporting the detection of self-oriented protection throughout adolescence, transitioning from local, memory-based processing in early adolescence to integrated hippocampus-prefrontal processing in late adolescence.

Our findings support that coordinated subcortical-cortical activity develops progressively throughout adolescence, with relevance for threat-safety discrimination. Younger adolescents appear to compensate for still-developing prefrontal systems by relying more heavily on hippocampal mechanisms when evaluating self-oriented protection cues. In the context of safety processing, the hippocampus may facilitate learning and memory retrieval about safety-relevant information before integrated hippocampal-prefrontal circuits take over these functions later in development. Importantly, Sastre et al. (2016) demonstrated that hippocampal activation during episodic retrieval is higher in children and decreases with age, which dovetails with prefrontal regions becoming more engaged to exert inhibitory control over the hippocampus, following a similar developmental trajectory to what we observed during safety evaluation. As adolescents mature, they transition from relying primarily on hippocampal processing to a more integrated circuit involving the vmPFC. This developmental pattern is consistent with previous research showing progressive strengthening of long-range connections between the hippocampus and prefrontal cortex throughout adolescence (Calabro et al., 2020; Murty et al., 2016). The development of these integrated circuits facilitates the retrieval and incorporation of relevant prior experiences and the formation of sophisticated schemas about threats and safety, both informing optimal behavioral responses to novel situations (Gluth et al., 2015; Spalding et al., 2015; van Kesteren et al., 2010; Voss et al., 2015). The importance of this circuitry for safety processing has been demonstrated in adults, with Tashjian et al. (2025) showing that the anterior vmPFC is specifically involved in processing self-oriented protection cues. Similarly, Milad et al. (2008) found that successful recall of extinction memory activates the vmPFC and hippocampus in concert, and Harrison et al. (2017) identified the vmPFC as critical for processing safety signals using a classical fear conditioning paradigm. In our task, participants needed to integrate information about their own protective capabilities with external threat information. The developmental shift toward greater reliance on hippocampal-vmPFC connectivity may reflect increasing capacity for this integration with age.

The transition to cortical-subcortical circuit functioning may create a period of vulnerability in older adolescents as they move away from established hippocampal mechanisms before fully developing prefrontal integration. The hippocampus stores episodic memories and contextual information, while the prefrontal cortex provides executive control, planning, and decision-making capabilities. Their connectivity allows past experiences (stored in the hippocampus) to directly inform complex decision-making processes (mediated by the PFC) (Preston & Eichenbaum, 2013; Murty et al., 2016). For threat-safety discrimination, the developmental vulnerability we observed in older adolescents likely reflects a transitional period when they have begun to switch to more sophisticated hippocampal-prefrontal circuits for safety judgments. Developing hippocampal-prefrontal circuits requires greater cognitive resources and coordination than either mature integrated networks or established single-region processing. Therefore, unlike younger adolescents who effectively use hippocampal processing alone, or adults with fully developed hippocampal-prefrontal connectivity, older adolescents may experience a temporary functional gap—they can no longer rely solely on hippocampal mechanisms but have not yet optimized the integration of prefrontal control systems. This creates a “developmental dip” in threat-safety discrimination accuracy during mid-to-late adolescence that may contribute to the increased risk-taking behaviors and vulnerability to anxiety observed during this period (Britton et al., 2013; Hur et al., 2025; Klein et al., 2024).

Interestingly, while we found age-related increases in hippocampal-ACC connectivity, similar to hippocampal-vmPFC connectivity, only the latter predicted threat-safety discrimination accuracy in our data. This may appear to contradict Meyer et al. (2019), who found that hippocampus-ACC connectivity, but not hippocampus-anterior vmPFC connectivity, was involved in threat inhibition via safety signals. However, we interpret our findings as extending rather than contradicting those of Meyer and colleagues. Specifically, hippocampus-anterior vmPFC connectivity is likely involved in safety processing when protection is perceived as independent from threat, whereas hippocampus-ACC connectivity might be more relevant for safety signals that directly inhibit threat processing (Wu et al., 2023; Jovanovic et al., 2013). Although we did not find a significant association between hippocampal-ACC connectivity and safety estimation in the current study, this proposed distinction between neural circuits that process safety independently of threat and those that modify threat directly may have important implications for clinical interventions. Therapeutic approaches can differentially target self-oriented safety processing or threat inhibition depending on the specific anxiety symptom profile, potentially increasing efficacy by addressing the distinct neural mechanisms underlying anxiety subtypes (Battaglia et al., 2022; Laing et al., 2022).

Results from the current study also clarify how competition between threat and safety cues is resolved within the visual cortex, demonstrating a preference for safety conferred via protection. Previous evidence suggests that the ventral visual networks play pivotal roles in tracking stimuli that are salient or highly relevant to behavior (Lang & Bradley, 2010; Wang et al., 2022) and supporting memory retrieval for the stimulus itself or similar experiences (Hofstetter et al., 2012). Among the most salient of all stimuli is threat, to which the sensory system exhibits bias for a survival response (Damaraju et al., 2009; Padmala & Pessoa, 2008; Stolarova et al., 2006). This preference for threat persists with the co-occurrence of threat and safety signals (Miskovic & Keil, 2013; Talmi et al., 2019). However, and contradicting previous studies, we found a preference in the visual system for safety compared to threat, which may be attributed to different constructs of safety. Building upon the conditioning framework, previous studies conceptualize safety as the absence or inverse of threat, and a threat-safety competition is evoked during safety acquisition, wherein the presence of a safety cue eliminates the significance of the threat cue. In the current study, safety is instead conferred in the presence of threat through protective weapons. These protection cues more closely resemble threat cues that are linked to sensory amplification—they are highly relevant and salient on their own, rather than having meaning for the relevance of an independent threat cue. In this case, sensory systems respond with increased activation to protection. This pattern aligns with predictive coding frameworks suggesting that when protection cues carry high informational value for survival outcomes, the visual cortex prioritizes their processing to optimize perceptual decision-making, contrasting with traditional threat-biased attention allocation (Mobbs et al, 2015; Price & Gavornik, 2022). The distinction between previous studies and the current one shows that actively increasing safety through positive cues reliably facilitates engagement of lower-tier sensory cortices, optimizing perception of safety and related decision-making.

Several limitations should be considered when interpreting our findings. First, while our sample size was adequate for detecting the reported effects, particularly with 7T fMRI, larger samples would provide greater opportunity for examining individual differences, and longitudinal inquiries would support interrogating developmental trajectories. Our paradigm focused on binary win/lose judgments rather than continuous ratings, the latter of which might have captured more subtle variations in threat-safety discrimination, and our use of behavioral responses rather than skin conductance response (SCR) as used in previous work. We used naturalistic threat and safety cues, presenting stimuli that are both perceptually threatening and modifying their safety value across matching probability spectrums. However, examining how different types of safety cues (e.g., social versus non-social) engage these neural systems across development could provide further insights into adolescent vulnerability and resilience.

By using 7T fMRI and examining both external threat and self-oriented protection cues, our study provides novel insights into the complex neurodevelopmental processes underlying safety evaluation during adolescence. Our findings reveal a developmental shift in the neural mechanisms supporting threat-safety discrimination during adolescence, transitioning from predominant reliance on hippocampal processing in early adolescence to integrated hippocampal-prefrontal processing in late adolescence. In addition, the visual cortex is involved in resolving competition between threat and safety cues, facilitating efficient identification and discrimination of safety-relevant information. These findings have implications for understanding adolescent risk-taking behavior and vulnerability to anxiety, potentially guiding the development of interventions aimed at strengthening threat-safety discrimination abilities during this critical developmental period. Interventions targeting threat-safety discrimination may be most effective when tailored to adolescents’ developmental stage, with younger adolescents potentially benefiting from memory-based approaches that leverage hippocampal processing, while older adolescents may require interventions that support the maturation of integrated prefrontal-subcortical circuits. Furthermore, the identified period of vulnerability in mid-to-late adolescence may represent a critical window for implementing preventive mental health strategies before maladaptive threat-safety patterns become entrenched.

## Glossary

Safety estimation: the cognitive process of evaluating the likelihood of safety (typically survival or absence of harm) when threats are present.
Threat-safety discrimination: the ability to differentiate between different threat and safety probabilities.
Safety prediction: the process of estimating safety likelihood based on incomplete information, specifically when only one component (either threat or protection) is known.
Safety integration: the process of combining multiple safety-relevant cues to form an updated, comprehensive safety assessment.

## CRediT Authorship Contribution Statement

**Yubing Zhang:** Conceptualization, Methodology, Software, Formal analysis, Investigation, Data Curation, Writing-Original Draft, Writing-Reviewing and Editing, Visualization. **Madeline Coates:** Investigation, Writing-Reviewing and Editing. **Marta I. Garrido:** Writing-Reviewing and Editing, Supervision. **Sarah M. Tashjian:** Conceptualization, Methodology, Software, Validation, Formal analysis, Resources, Data Curation, Writing-Reviewing and Editing, Visualization, Supervision, Project administration, Funding acquisition.

## Declaration of Competing Interest

The authors declare that they have no known competing financial interests or personal relationships that could have appeared to influence the work reported in this paper.

## Funding Sources

This work was supported by a National Health and Medical Research Council (NHMRC) Investigator Grant (2033400) and a Brain and Behavior Research Foundation Young Investigator Grant to SMT (30788). The funders played no role in study design, data collection or analysis, decision to publish, or preparation of the manuscript.

## Acknowledgements

We extend our deepest gratitude to the adolescents who participated in this study as well as their parents/guardians. Thank you to Heidi Meyer, PhD, and Dylan Gee, PhD, for their assistance with defining our hippocampal ROI.

## Code and Data Availability

Task code and behavioral data are available through the Open Science Framework (OSF) (https://doi.org/10.17605/OSF.IO/V3KPX). Neuroimaging data are available through the Science Data Bank (https://www.scidb.cn/en/anonymous/QmpZUk52).

## Notes

### Competing Interest Statement

The authors have declared no competing interest.

